# Electromyographic analysis of the suprahyoid muscles in infants based on the lingual fraenulum attachment during breastfeeding

**DOI:** 10.1101/488437

**Authors:** Ellia Christinne de Lima França, Lucas Carvalho Aragão Albuquerque, Roberta Lopes de Castro Martinelli, Ilda Machado Fiuza Gonçalves, Cejana Baiocchi Souza, Maria Alves Barbosa

## Abstract

**Introduction:** Muscle electrical activity analysis can aid in the identification of oral motor dysfunctions such as those resulting from altered lingual fraenulum which, in turn, impairs feeding. We aimed to analyse suprahyoid muscle electrical activity of infants based on lingual fraenulum attachment to the sublingual (ventral) aspect of the tongue and floor of the mouth, during breastfeeding.

**Methods and Results:** We studied full-term infants of both genders aged between 1–4 months. Lingual fraenulum evaluation and surface suprahyoid muscle electromyography was performed during breastfeeding. Mean muscle activities were recorded in microvolts and converted into percent values (normalisation) of the reference value. Associations between root mean square and independent variables were tested by one-way analysis of variance and Student’s t-test with significance level of 5% and test power of 95%.

We evaluated 235 infants while breastfeeding. The lingual fraenulum was commonly attached to the tongue’s ventral aspect between middle third and apex, and on the mouth floor visible from the lower alveolar ridge. Lower muscle activity was observed with lingual fraenulum attached to apex/lower alveolar ridge, followed by attachment to middle third/lower alveolar ridge, and between middle third and apex/lower alveolar ridge. Highest activity observed in Infants with attachment to middle third/sublingual caruncles, had a thin lingual fraenulum, performed several sucks followed by short pauses, showed coordination between swallowing, sucking, and breathing, did not “bite” nipple, and showed no tongue snapping nor stress.

**Conclusion:** Greater suprahyoid muscle activity during breastfeeding was observed with lingual fraenulum attachment to middle third of the tongue/sublingual caruncles, showed coordination between swallowing, sucking, and breathing. Surface electromyography is effective in diagnosing lingual fraenulum alterations, whose attachment point raises doubts as restriction of tongue mobility. This technique identifies possible oral motor dysfunctions, enables direct therapeutic interventions and early intervention, and prevents feeding and communication alterations.

## Introduction

Breastfeeding is considered essential for the promotion and protection of children’s health due to the nutritional and immunological properties of breast milk, which protect them from respiratory diseases, infectious [1], and diarrhea [2]. In addition, breastfeeding is considered important for the adequate development of the stomatognathic system, because the removal of breast milk involves intense muscle activity [3].

During sucking in the womb, the suprahyoid muscles (digastric, mylohyoid, geniohyoid, and stylohyoid) that effectively participate in the movement and stabilisation of the mandible and tongue movement [4,5].

The correct movement of the tongue during breastfeeding promotes an adequate fit between the infant’s mouth and the mother’s nipple, compressing the nipple against the hard palate and favouring the removal of the milk due to the vacuum created in the oral cavity by the raising and lowering movement of the tongue [6–9].

In the inferior surface of the tongue is located the lingual fraenulum and is considered a median fold of tunica mucosa that connects the tongue to the floor of the mouth and allows the free movement of its anterior part [10].

After apoptosis, the remaining residual embryonic tissue (lingual fraenulum) may limit tongue movements to varying degrees. This congenital oral anomaly is referred to as ankyloglossia [11] and may cause a reduced mouth opening, imprecision and restriction of isolated tongue movements, heart-shaped tip or downward protrusion [12], tongue resting in the floor of the mouth, and difficulties in sucking, chewing, swallowing, and speech functions, among others [11–15].

Oral dysfunctions caused by an altered lingual fraenulum may compromise breastfeeding by causing discomfort and pain to mothers, ineffective emptying of the breast [11], poor weight gain, and/or early weaning [10].

Studies that evaluated the lingual fraenulum by quantitative and objective methods are scarce and tried to determine whether the anatomical findings could compromise tongue movement, and consequently the oral functions [6]. Such methods enable safe diagnoses to guide therapeutical procedures for the continuity of breastfeeding.

Research on the use of surface electromyography (EMG), which is a method for recording variations in muscle electrical activity during contraction [16,17], are also scarce, especially those evaluating the sucking function in infants [5,18–26]. EMG is considered an easy, fast, low-cost, safe, and non-invasive procedure that can provide important information about muscle functions [5,16–18,23–26], which can be used to diagnose oral-motor dysfunction accurately [23,24].

Some studies have evaluated the activity of orofacial muscles using surface electromyography during breastfeeding and other feeding methods [5,18–24,26] but, to the best of our knowledge, no study analysed the activity of the suprahyoid muscles in newborns and infants during the sucking function in breastfeeding based on the lingual fraenulum attachment.

We decided to analyse the suprahyoid muscles (digastric, mylohyoid, geniohyoid, and stylohyoid), because they directly participate in the sucking function, act in mandible movement and stabilisation, and participate in tongue movement, and, because of their location, makes it possible to attach the electrodes [5].

There is still no established pattern for the electrical activity of the suprahyoid muscles involved in the sucking function in infants based on the lingual fraenulum attachment to the tongue and the floor of the mouth. In this context, we note the importance of in-depth studies on the subject that may favour the understanding of muscle activity, and aid in the identification of possible oral motor dysfunctions and feeding efficiency. These studies may also contribute to the planning and implementation of actions in health services that would enable breastfeeding and minimise the impact and consequences related to early weaning, which would benefit child development.

The present study’s aim was to analyse the electrical activity of the suprahyoid muscles in infants, based on the lingual fraenulum attachment to the tongue and the floor of the mouth, during the sucking function in breastfeeding.

## Materials and Methods

### Study Design

This is an observational, analytical, cross-sectional study. The Research Ethics Committee approved the study under number 705.229 on June 30, 2014. This study was conducted at a University in the central region of Brazil. All adults responsible for the infants participating in this study received and signed an Informed Consent Form and a Consent Form for the Participation of the Person as a Subject, respecting Resolution 466/2012.

### Participants

Infants of both genders aged between 1 and 4 months had their lingual fraenulum evaluated from March 2015 to December 2016. Infants with anatomical and physiological changes on the face, pre-term and post-term neurological impairment, weight below 2500 g, or exclusively bottle-fed were excluded.

### Procedures

The independent variables analysed were: age in days at the examination, gender, clinical history (CH), lingual fraenulum (visualisation is possible, visualisation is not possible, or visualisation is only possible with manoeuvre), lingual fraenulum thickness (thin or thick), lingual fraenulum attachment to the tongue (middle third, between the middle third and the apex, or at the apex) and the floor of the mouth (visible from the sublingual caruncle or lower alveolar ridge), non-nutritive sucking pattern – NNS (adequate: forward positioning of the tongue, coordinated movements and efficient sucking, inadequate: limited forward positioning of the tongue, uncoordinated movements and delayed onset of sucking) and nutritive sucking - NS (rhythm: several sucks followed by short pauses or few sucks followed by long pauses); coordination: adequate - balance between food efficiency and sucking functions, swallowing and breathing, no signs of stress or inadequate - cough, choking, dyspnoea, regurgitation, hiccups, swallowing noises; “Bites” the nipple: yes or no; tongue snapping during sucking: yes or no).

The dependant variable analysed was muscle electrical activity (MEA) defined by the mean of the potential action of the motor units of a muscle group, obtained from the electromyographic signal, and expressed in root meant square (RMS) in microvolts (μV) and later converted to percentage (%).

We used the “Protocol for evaluation of lingual fraenulum in infants” proposed by Martinelli [27] for the quantitative evaluation of lingual fraenulum.

The parents were advised at the time of the examination not to feed the babies for at least two hours before the evaluation.

The first stage of the protocol was the data collection of the clinical history, including the following items: date of the examination, full name, gender, birth date, age, address, telephone number, parents’ name, family members with lingual fraenulum alterations, data on the infant’s current general health, time between feedings, data on whether the infant gets tired while sucking, and data on whether the infant progressively releases the nipple or bites the nipple.

The second stage of the protocol included an anatomical and functional evaluation to observe the general aspects of the lingual fraenulum, a nutritive sucking evaluation during breastfeeding, to investigate tongue movements and positioning in the oral cavity, through electromyographic.

During the anatomical and functional evaluation, we observed the posture of the lips at rest (closed, semi-open, or open), the tendency of tongue positioning during crying (raised, midline, midline with lateral elevation, or downward tongue tip with lateral elevation), and the shape of the tip of the tongue when raised during crying or a lifting manoeuvre (rounded, slight crevice at apex, or heart-shaped). Through the elevation of the lateral margins of the tongue by the gloved right and left index fingers of the evaluator, it was observed whether lingual fraenulum could be visualised or not and if it was necessary to visualised with specific manoeuvre. If the lingual fraenulum could be visualized, we determined: whether it was thin or thick; whether the lingual fraenulum was attached to the middle third of the tongue, between the middle third and the apex, or to the apex; and whether the attachment to the floor of the mouth was visible from the sublingual caruncles (opening of the ducts of the right and left submandibular glands) or from the lower alveolar ridge.

The non-nutritive sucking was evaluated by introducing the gloved minimum finger into the baby’s mouth during sucking for 2 minutes, and tongue movement was observed (adequate: tongue anterioration, coordinated movements and efficient suctioning; inadequate: limited tongue anterioration, uncoordinated movements and delayed onset of suction).

The evaluation of nutritive sucking was made along with surface electromyography. We observed: the sucking rhythm (counting the number of sucks in three sucking groups, which were separated by pause time, and then taking the mean); pause time (considering the duration of the pauses between three sucking groups, and then taking the mean); coordination between sucking/swallowing/breathing and classified into adequate or inadequate; and whether the infant bit the nipple and whether the infant had tongue snapping during nutritional sucking.

Evaluation of the electrical activity of the suprahyoid muscles (MEA) included all infants and was performed during sucking of the mother’s breast. We used the MIOTOOL 200 device (manufactured by Miotec Equipamentos Biomédicos Ltda. - ME, Porto Alegre, Brazil) that was comprised of four channels, had a 7.2 V 1.700 Ma NiMH rechargeable battery, was operated in isolation from the utility system, and was connected to a Sony Vaio^®^ notebook. The electromyographic signal was processed through a data acquisition system that allowed for the selection of 8 independent gains per channel, which was used to a gain of 1000; a low pass filter of 20 Hz; a high pass filter of 500 Hz; two SDS500 sensors connected to clamp sensors; a reference cable (earth); and a calibrator (MIOTEC^®^) were used.

The records were made in a quiet place with natural light and ambient temperature [28]. We used disposable unipolar surface electrodes (Meditrace^®^, Infant Model, manufactured by Tyco/Kendall-USA, imported by Lamedid Comercial e Serviços Ltda, Barueri - SP - Brazil São Paulo, Brazil) with a material made of silver-silver chloride (Ag-AgCl), adhesive, and conductive solid gel (hydrogel) that were responsible for capturing and conducting the signal of the EMG. Subsequently, the skin was cleaned with gauze soaked in 70° alcohol to remove oil or any material that interfered with the signal capture [29,30].

The electrode positioning followed a standard procedure that started with the reference electrode or “earth”, followed by the attachment of the electrodes to the suprahyoid muscles. The reference electrode, which was placed in the frontal bone (forehead), minimises interferences from external electrical noises [17,29].

The other electrodes were placed in a bipolar configuration in the suprahyoid muscles with a minimum distance of 10 mm between them [30]. The evaluator stimulated non-nutritive sucking by introducing a flavoured (milk breast) finger for five seconds to locate this region and, by this manoeuvre, it was possible to palpate the muscles.

After attaching the electrodes to the skin of the infant, the clamp sensors were placed following the same order as the electrode attachment [17,29]. After completing this procedure, the configuration, channel enablement in the software, and subsequent calibration were performed. The three unused channels were disabled.

The mother was seated comfortably in a chair with back support with her baby on her lap. Before the evaluation, she received the necessary guidelines on the clinical examination, evaluation of nutritive sucking, and electromyographic examination.

The mother received instructions about her positioning: seated with feet on the ground; and about the positioning of the infant: supported and aligned with the head and the spine straight, belly facing the body of the mother, face towards the mother’s breast, and mouth towards the areola and nipple to catch the breast.

Subsequently, the activity of the suprahyoid muscles during breastfeeding was recorded for 3 minutes (Fig 1).

**Figure 1.**
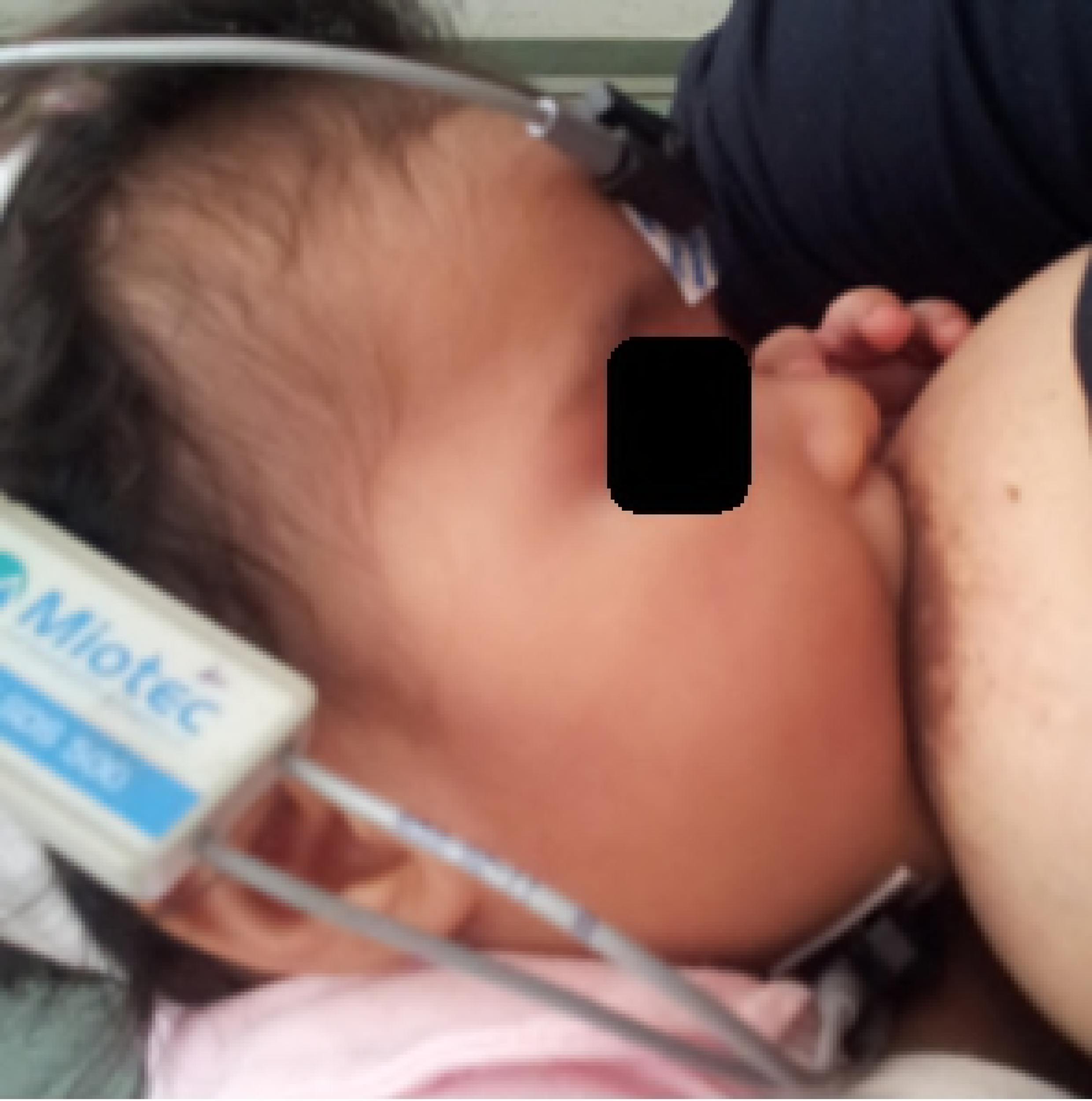
Electromyographic evaluation. Electrodes attached to bone (forehead) and submandibular (suprahyoid muscles) regions during sucking in breastfeeding

The infants whose results showed interference of the lingual fraenulum in the tongue movements and/or with reduced activity of the suprahyoid muscles were referred to basic health units with a speech-language pathology report of the lingual fraenulum. After scheduling a consultation with the paediatrician, the infant was referred to the paediatric dentistry service for assessment and definition of conduct.

The data collected in this study were archived in a confidential location and will be incinerated after 5 years.

### Electromyographic analysis

The Miograph 2.0 (MIOTEC ^®^, São Paulo, Brazil) software was used for presenting and interpreting the electromyographic signal. The numerical data was expressed in *root mean square* (RMS), which represents the square root result of the mean square of the instantaneous amplitudes of the signal of the recorded electromyographic trace whose unit is expressed in microvolts (μV).

In order to select the best signals, the best configurations that presented the least noise and the most symmetrical and connected histogram to the signal were considered.

The means recorded in μV were transformed into percent values of the reference value (normalisation by the maximum peak) for each subject. The formula for calculating the percentage, according to the recommendations of the International Society of Electrophysiology and Kinesiology (ISEK) [30], was (X/Y) × 100 where X = MEA (muscular electrical activity) mean in the requested task (μV) and Y = reference value corresponding to the mean of the MEA in PICO (μV). Thus, the highest value of the electromyographic signal of the suprahyoid muscles was identified for 3 seconds of breastfeeding. The maximum peak was considered 100% of activity and the mean activity during 3 seconds of the breastfeeding was considered “X” [17,29,31].

The normalisation technique is essential so that the surface electromyography signal can be compared in different studies, muscles, and participants after being analysed. This technique is considered a prerequisite for any comparative analysis of EMG signals [28,31].

### Statistical analysis

The data were entered into IBM SPSS Statistics v. 23 (IBM, Inc, Chicago, Illinois, USA) and subjected to a descriptive, inferential, and analytical statistical analysis using frequencies, central tendency, and dispersion measure. A test power of 95% was used, and type 1 error was set at 5%.

The muscular electrical activity found in RMS corresponded to the dependant variable, which had a normal distribution (Kolmogorov-Smirnov p = 0.113), considering both groups. The independent variables were: age in days at the electromyographic examination, gender, feeding type, lingual fraenulum attachment type, and nutritive sucking pattern. The association between the dependant variable RMS and the independent variable age was tested by a one-factor Analysis of Variance (ANOVA), which allows for the comparison of continuous variables at a single time. The association between the dependant variable RMS and other independent variables was tested by the Student’s t-test, which allows for the comparison of two samples using their means for statistical inference.

## Results

Two hundred fifty-one infants that met the inclusion criteria underwent surface electromyography between February 2015 to November 2016. Eleven infants were excluded because their records showed an electromyographic signal with interference, and four infants were excluded due to a failure in the capture and recording of the electromyographic signal at the time of evaluation that resulted in a final sample size of 235 infants.

The highest age range at the day of the examination was 31 to 60 days, corresponding to 37.0% (n = 87) of the infants, and was followed by 61 to 90 days, corresponding to 26.8% (n = 63) of the infants. Only 5.1% (n = 12) of the infants were aged between 121 and 149 days.

Most of the infants were female, corresponding to 51.9% (n = 122) of the total sample, and 48.1% (n = 113) were male.

Exclusive breastfeeding was the feeding type most used, corresponding to 176 infants (74.9%), and was followed by breastfeeding/bottle, corresponding to 59 babies (25.1%). The justifications reported by the mothers were the child crying a lot, irritation, being hungry, weak, insufficient breast milk and not breastfeeding gain weight. All of the infants were assessed by surface electromyography during sucking of the maternal breast.

Of the 235 infants evaluated, 197 (83.8%) did not have complaints of difficulties in breastfeeding in their clinical history; 172 (73.1%) had a lingual fraenulum that could be visualised without manoeuvre; in 63 (27.3%) of the infants presented lingual frenulum covered by a mucosal curtain, being necessary maneuver to visualize it; 215 (91.4%) had a thin lingual fraenulum and 20 (9.3%) thick lingual fraenulum. The infants were divided into six groups according to the location of the fraenulum attachment on the tongue and floor of the mouth: 53 (22.5%) had the lingual fraenulum attached of the middle third on the tongue and visible from the sublingual caruncles (group 1); 35 (14.8%) had the lingual fraenulum attached to the middle third/lower alveolar ridge (group 2); 28 (11.9%) had attached between the middle third and apex, and visible from the sublingual caruncles (group 3); 102 (43.4%) had the fraenulum attached between the middle third and the apex and visible from the lower alveolar ridge (group 4); 17 (7.2%) had the fraenulum attached to the apex and visible from the lower alveolar ridge (group 5); and no infants with the lingual fraenulum attached in the apex on the tongue and visible from the caruncles were identified (group 6).

Of the 235 infants evaluated, 146 (62,1%) presented adequate nutritive sucking.

For the statistical analysis associating the dependant variable of muscular electrical activity value with the other variables, were excluded the data referring to items 1, 2, and 3 of the anatomical and functional evaluation, because they are not directly related to the anatomy of the lingual fraenulum and non-nutritive sucking (NNS), not to be the object of study of this work.

The data from the protocol for lingual fraenulum evaluation in infants were analysed and the results from the anatomical and functional evaluation part I item 4 (which analysed the lingual fraenulum thickness and fraenulum attachment to the sublingual aspect and the floor of the mouth), and the NS pattern (which comprised the rhythm, coordination, nipple biting, and tongue snapping during sucking) were aggregated.

There was no statistically significant difference (Table 1) when electrical activity of the suprahyoid muscles was associated with the age of the infants (p = 1 0.368) (ANOVA), gender (p = 0.136), and feeding type (p = 0.689) (T-test).

**Table 1.**
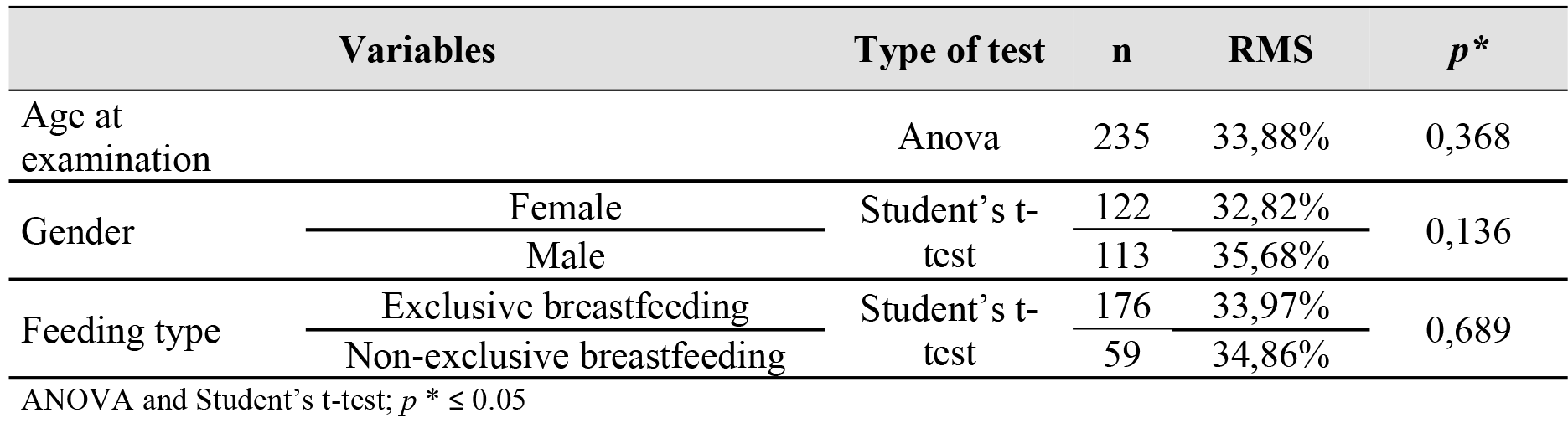
Association between the characteristics of infants (age in days at the examination, gender, and feeding type) and muscular electrical activity (MEA).

We observed (Table 2) a greater electrical activity in the muscle during breastfeeding in the infants with their lingual fraenulum attached to the middle third and visible from the sublingual caruncle (40.69%), and lower electrical activity in the muscle in the infants with their lingual fraenulum attached to the apex of the tongue and visible from the lower alveolar ridge (29.94%).

**Table 2.** Mean activity of the suprahyoid muscles expressed in RMS of infants based on the lingual fraenulum attachment to the tongue and the floor of the mouth during breastfeeding.

| Group | Location of lingual fraenulum attachment |  | n | RMS | DP |
| --- | --- | --- | --- | --- | --- |
|  | Sublingual | Floor of the mouth |  |  |  |
| 1 | middle third | sublingual caruncles | 53 | 40,69% | 15,22 |
| 2 | middle third | lower alveolar ridge | 35 | 31,13% | 11,69 |
| 3 | middle third/apex | sublingual caruncles | 28 | 36,91% | 12,94 |
| 4 | middle third/apex | lower alveolar ridge | 102 | 31,84% | 13,82 |
| 5 | apex | lower alveolar ridge | 17 | 29,94% | 19,45 |

Student's t-test; SD - Standard Deviation; RMS - *root mean square*

A statistically significant difference was found when comparing the electrical activity of the muscle and the location of the lingual fraenulum attachment on the tongue and the floor of the mouth of infants, according to the division of the groups. A significant difference was found between group 1 (middle third/sublingual caruncles) and group 2 (middle third/lower alveolar ridge) (p = 0.002), between group 1 (middle third / sublingual caruncles) and group 4 (middle third and apex/lower alveolar ridge) (p = 0.001), and between group 1 (middle third/sublingual caruncles) and group 5 (apex/lower alveolar ridge) (p = 0.021) (Table 3).

**Table 3.** Comparison of the electrical activity in the suprahyoid muscles between groups expressed in RMS of infants during breastfeeding based on the lingual fraenulum attachment to the tongue and floor of the mouth.

| Group | Location of lingual fraenulum attachment | | n | RMS | $p^*$ |
| --- | --- | --- | --- | --- | --- |
|  | Sublingual | Floor of the mouth |  |  |  |
| 1 and 2 | middle third | sublingual caruncles | 53 | 40,69% | 0,002 |
|  | middle third | lower alveolar ridge | 35 | 31,13% |  |
| 1 and 4 | middle third | sublingual caruncles | 53 | 40,69% | 0,001 |
|  | middle third / apex | lower alveolar ridge | 102 | 31,84% |  |
| 1 and 5 | middle third | sublingual caruncles | 53 | 40,69% | 0,021 |
|  | apex | lower alveolar ridge | 17 | 29,94% |  |

Student's t-test; $p^* \leq 0.050$ ; RMS - *root mean square*

Considering the statistically significant differences found in the association between the groups, as described in Table 3, the electrical activity of the muscle was compared to the location of the fraenulum attachment in the tongue and floor of the mouth, lingual fraenulum thickness (thin or thick), and the pattern of nutritive sucking (rhythm, coordination, bit the nipple, and tongue snapping). There was a statistically significant difference, with a higher mean percentage of the electrical activity of the suprahyoid muscle, during breastfeeding in infants with the lingual fraenulum attached to the middle third/sublingual caruncles (group 1), who had a thin lingual fraenulum (p = 0.002), with several sucks and short pauses during NS (p = 0.003), coordination between sucking, swallowing and breathing functions, without signs of stress (p = 0.001), absence of bite in the nipple (p = 0.005), and absence of tongue snapping (p = 0.001) when compared to infants with the lingual fraenulum attached to the middle third/lower alveolar ridge (group 2) (Table 4).

**Table 4.** Comparison of the electrical activity in suprahyoid muscles between groups 1 and 2 expressed in RMS of infants according to thickness, lingual fraenulum attached in the tongue and floor of the mouth, and nutritive sucking pattern.

| Variables |  | Lingual fraenum attachment/floor of the mouth | *n* | RMS | *p* * |
| --- | --- | --- | --- | --- | --- |
| Fraenum Thickness | Thin | G1-MT / SC | 47 | 41,15% | 0,002 |
|  |  | G2 - MT / IAC | 35 | 31,12% |  |
|  | Thick | G1-MT / SC | 6 | 37,06% | 0 |
|  |  | G2 - MT / IAC | 0 | 0,00% |  |
| Rhythm | Several sucks / short breaks | G1-MT / SC | 46 | 41,72% | 0,003 |
|  |  | G2 - MT / IAC | 32 | 31,68% |  |
|  | Few sucks / long breaks | G1-MT / SC | 7 | 33,92% | 0,31 |
|  |  | G2 - MT / IAC | 3 | 25,24% |  |
| Coordination | Proper | G1-MT / SC | 51 | 41,38% | 0,001 |
|  |  | G2 - MT / IAC | 34 | 31,13% |  |
|  | Inadequate | G1-MT / SC | 2 | 23,08% | 0,61 |
|  |  | G2 - MT / IAC | 1 | 31,01% |  |
| Bite the nipple | Not | G1-MT / SC | 48 | 41,77% | 0,005 |
|  |  | G2 - MT / IAC | 26 | 31,85% |  |
|  | Yes | G1-MT / SC | 5 | 30,33% | 0,84 |
|  |  | G2 - MT / IAC | 9 | 29,03% |  |
| Tongue snapping | Not | G1-MT / SC | 36 | 42,55% | 0,001 |
|  |  | G2 - MT / IAC | 18 | 27,48% |  |
|  | Yes | G1-MT / SC | 17 | 36,75% | 0,71 |
|  |  | G2 - MT / IAC | 17 | 34,98% |  |

Student's t-test; $p^* \leq 0.050$ ; RMS - *root mean square*
Group 1- Middle third / sublingual caruncles ( $n = 53$ )
Group 2 - Middle third / lower alveolar ridge ( $n = 35$ )

A higher mean electrical activity of the suprahyoid muscles was observed during nutritive sucking in infants with the lingual fraenulum attached to the middle third/sublingual caruncle (group 1), who had a thin lingual fraenulum (p = 0.001), with coordination between the sucking (p = 0.003), swallowing and breathing functions, without signs of stress (p = 0.001), absence of bite at the nipple (p = 0.001), and absence of tongue snapping (p = 0.001) when compared to infants with the lingual fraenulum attached to the middle third and apex/lower alveolar ridge (group 4) (Table 5).

**Table 5.** Comparison of the electrical activity in the suprahyoid muscles between groups 1 and 4 expressed in RMS of infants according to thickness, fraenulum attachment to the tongue and floor of the mouth, and nutritive sucking pattern.

| Variables |  | Lingual fraenum attachment/floor of the mouth | n | RMS | p* |
| --- | --- | --- | --- | --- | --- |
| Fraenum Thickness | Thin | G1 - MT / SC | 47 | 41,15% | 0,001 |
|  |  | G4 - MTA / IAC | 94 | 31,57% |  |
|  | Thick | G1 - MT / SC | 6 | 37,06% | 0,8 |
|  |  | G4 - MTA / IAC | 8 | 35,02% |  |
| Rhythm | Several sucks / short breaks | G1 - MT / SC | 46 | 41,72% | 0,003 |
|  |  | G4 - MTA / IAC | 66 | 33,19% |  |
|  | Few sucks / long breaks | G1 - MT / SC | 7 | 33,92% | 0,4 |
|  |  | G4 - MTA / IAC | 36 | 29,37% |  |
| Coordination | Proper | G1 - MT / SC | 51 | 41,38% | 0,001 |
|  |  | G4 - MTA / IAC | 91 | 32,71% |  |
|  | Inadequate | G1 - MT / SC | 2 | 23,08% | 0,86 |
|  |  | G4 - MTA / IAC | 11 | 24,68% |  |
| Bite the nipple | Not | G1 - MT / SC | 48 | 41,77% | 0,001 |
|  |  | G4 - MTA / IAC | 78 | 32,34% |  |
|  | Yes | G1 - MT / SC | 5 | 30,33% | 0,98 |
|  |  | G4 - MTA / IAC | 24 | 30,23% |  |
| Tongue snapping | Not | G1 - MT / SC | 36 | 42,55% | 0,001 |
|  |  | G4 - MTA / IAC | 34 | 29,30% |  |
|  | Yes | G1 - MT / SC | 17 | 36,75% | 0,34 |
|  |  | G4 - MTA / IAC | 68 | 33,12% |  |

Student's t-test; $p \leq 0.050$ ; RMS - root mean square
Group 1- Middle third / sublingual caruncles (n = 53)
Group 4 – Middle third and Apex / lower alveolar ridge (n = 102)

A statistically significant difference was observed with a higher mean percentage of the electrical activity of the suprahyoid muscles during breastfeeding in infants with the lingual fraenulum attached to the middle third/sublingual caruncles (group 1); who had a thin lingual fraenulum (p = 0.02); a NS pattern with coordination between sucking, swallowing, and breathing functions; without signs of stress (p = 0. 005); absence of bite at the nipple (p = 0.03); and absence of tongue snapping (p = 0.006) when compared to infants with the lingual fraenulum attached to the apex/lower alveolar ridge (group 5) (Table 6).

**Table 6.**
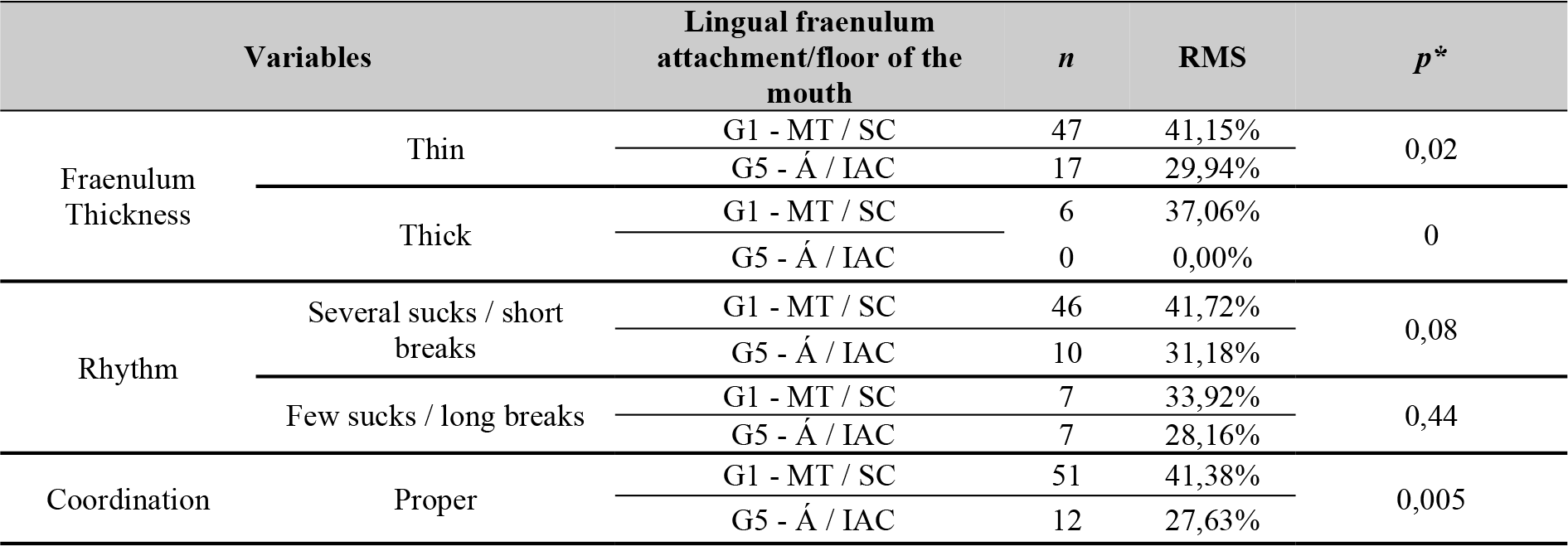
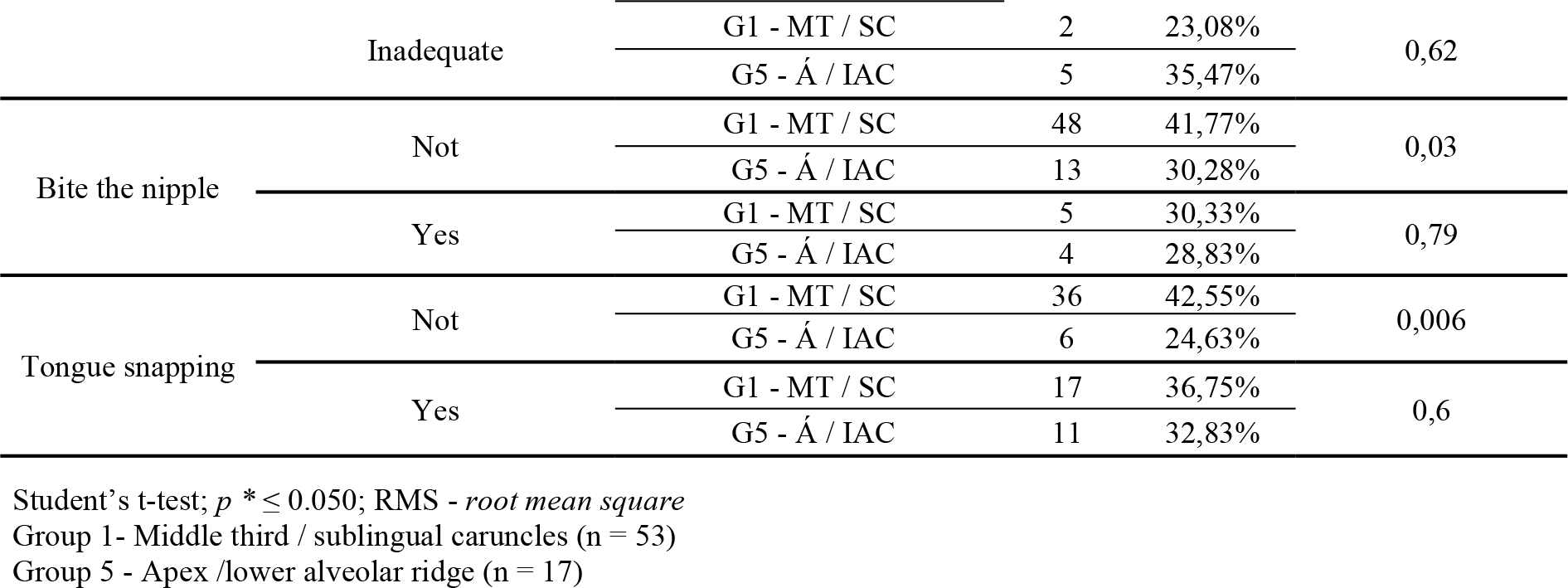
Comparison of the electrical activity in the suprahyoid muscles between groups 1 and 5 expressed in RMS of the infants according to thickness, lingual fraenulum attachment to the tongue and floor of the mouth, and nutritive sucking pattern.

| Variables |  | Lingual fraenum attachment/floor of the mouth | n | RMS | p* |
| --- | --- | --- | --- | --- | --- |
| Fraenum Thickness | Thin | G1 - MT / SC | 47 | 41,15% | 0,02 |
|  |  | G5 - Á / IAC | 17 | 29,94% |  |
|  | Thick | G1 - MT / SC | 6 | 37,06% | 0 |
|  |  | G5 - Á / IAC | 0 | 0,00% |  |
| Rhythm | Several sucks / short breaks | G1 - MT / SC | 46 | 41,72% | 0,08 |
|  |  | G5 - Á / IAC | 10 | 31,18% |  |
|  | Few sucks / long breaks | G1 - MT / SC | 7 | 33,92% | 0,44 |
|  |  | G5 - Á / IAC | 7 | 28,16% |  |
| Coordination | Proper | G1 - MT / SC | 51 | 41,38% | 0,005 |
|  |  | G5 - Á / IAC | 12 | 27,63% |  |
|  | Inadequate | G1 - MT / SC | 2 | 23,08% | 0,62 |
|  |  | G5 - Á / IAC | 5 | 35,47% |  |
| Bite the nipple | Not | G1 - MT / SC | 48 | 41,77% | 0,03 |
|  |  | G5 - Á / IAC | 13 | 30,28% |  |
|  | Yes | G1 - MT / SC | 5 | 30,33% | 0,79 |
|  |  | G5 - Á / IAC | 4 | 28,83% |  |
| Tongue snapping | Not | G1 - MT / SC | 36 | 42,55% | 0,006 |
|  |  | G5 - Á / IAC | 6 | 24,63% |  |
|  | Yes | G1 - MT / SC | 17 | 36,75% | 0,6 |
|  |  | G5 - Á / IAC | 11 | 32,83% |  |

Student's t-test; $p \leq 0.050$ ; RMS - *root mean square*
Group 1- Middle third / sublingual caruncles (n = 53)
Group 5 - Apex /lower alveolar ridge (n = 17)

## Discussion

The age group of 1 to 4 months analysed in the present study is the same as other studies that used electromyographic evaluation in infants [18–23,26]. Most of the infants were aged between 31 and 60 days, which favours the promotion of breastfeeding and an early diagnosis of the lingual fraenulum in the population attended, preventing eventual difficulties in feeding and speech [10,12–15].

Most of the infants were exclusively breastfed. Of the 235 infants included, 57 were already using a bottle to complement their feeding. The main reasons reported by the mothers for the bottle complementation included the child was crying a lot, irritated, hungry, weak, not gaining weight, or the mother was producing insufficient breast milk. These data agree with the literature. A previous study showed that the main reasons for early weaning included the mothers’ lack of knowledge about breastfeeding benefits, care and clinical management, and functional and anatomical problems related to the mothers and infants [32]. Another previous study identified some of the main difficulties in breastfeeding related to a lingual fraenulum alteration, which included increased feeding time, low breast milk intake, and poor weight gain [11,14,15,33], favouring early weaning [11,14,15,33–35].

All the infants that participated in this study were referred from public maternity hospitals that are regulated by the Brazilian Unified Health System (Sistema Único de Saúde - SUS). Currently, in the municipality of Goiânia in the State of Goiás, four maternity hospitals are qualified by the Brazilian Ministry of Health, receive financial incentives, and are included in the Baby-Friendly Hospital Initiative programme [36]. This support may have favoured exclusive breastfeeding in the infants of this study, since the main objectives of this programme are protection, promotion, and support of breastfeeding [37]. A previous study revealed [37,38] that infants born in maternities that do not participate in the Baby-Friendly Hospital Initiative were more likely to have early weaning.

In the anatomical and functional evaluation, we observed that 27.3% of the infants had their lingual fraenulum covered by a mucosal curtain, which was visualized through a manoeuvre, and 9.3% had a thick lingual frenulum. These findings agree with a previous study [15] that evaluated 100 healthy, full-term, 30-day-old infants. The authors in that study found the incidence of a lingual fraenulum covered by mucosal curtain to be 29%. Another study [39] performed with 1,084 healthy infants, found that 35% of these had posterior lingual fraenulum and that this type of lingual fraenulum did not interfere with breastfeeding. An experimental study [40], performed with 1,715 healthy infants, found the occurrence of posterior or submucosal lingual frenulum in 558 (32.54%) infants, as well as the efficiency of the maneuver used to visualize this lingual frenulum. There was a higher frequency (43.4%) of lingual fraenulum attachment to the tongue between the middle third and the apex and visible from the lower alveolar ridge. These results agree with the literature: a retrospective study that analysed 165 protocols for assessing the lingual fraenulum of full-term babies aged between 1 and 4 months that found a prevalence of 32.2% of the lingual fraenulum attached between the middle third and the apex/ lower alveolar ridge [39]. Another previous longitudinal study, which evaluated the anatomical characteristics of the lingual fraenulum of 71 full-term infants, found that 27 (38%) had their lingual frenulum attached between the middle third and the apex at the 1^st^, 6^th^, and 12^th^ month of life, and 42 (59.1%) infants had their lingual frenulum attached in the lower alveolar ridge [15].

In the present study, there was no significant difference between the electrical activity of the suprahyoid muscles during nutritional sucking and the infants’ age in days (p = 0.368), as opposed to the literature findings. A previous study [22] that investigated the development of sucking of full-term infants through electromyography during breastfeeding found that the activity of the suprahyoid muscles was higher with increasing age and that the activity of this muscle in the active movements of the tongue and mandible lowering favours an increase of sucking force in breastfeeding. Other previous studies that analysed the submandibular muscles in breastfeeding through ultrasonography found that the sucking pattern does not differ in infants aged between 1 to 4 months, indicating an early development of the coordination between sucking, swallowing, and breathing and that this may vary depending on the milk flow at the time of ejection and the adaptive capacity of infants during breastfeeding [8,9]

There was no significant difference between electrical activity of the muscle during sucking and gender (p = 0.136). Several previous studies [22,23,26] that analysed the activity of orofacial muscles of full-term infants and premature infants during different feeding methods did not find a correlation between the electrical activity of the studied muscles and gender. Although 59 (25.1%) of the infants analysed in the present study used complementary feeding with a bottle at home, the results did not have a significant difference when compared to feeding type (p = 0.689). This result may be because all electromyography tests were performed during sucking of the breast. It should be emphasised that the fact that mothers supplement the feeding of their babies at night with a bottle did not influence the electrical activity of the muscle. Some previous studies indicated that the use of bottle feeding favours the participation of the mental and buccinator muscles and causes a hypofunction of the masseter [18–20,23], temporal, pterygoid, tongue, and lip muscles [18].

In the present study, we found lower electrical activity of the suprahyoid muscles during sucking of the breast in infants with their fraenulum attached of the middle third of the tongue, between the middle third and the apex, and to the apex and visible from the lower alveolar ridge, regardless of age. These results indicate that the attachment of the lingual fraenulum on the mouth floor visible from the lower alveolar ridge, appears to interfere more with the tongue forward when compared to the fraenulum attached of the middle third of the tongue and middle third apex.

These results provide important data for the differential diagnosis of the lingual fraenulum, because they identify anatomical characteristics that may reduce muscle activity due to the restriction of tongue-tip movement during sucking in infants that could impair breastfeeding. A previous study [6] that assessed infants with persistent difficulties in breastfeeding through submandibular ultrasonography during breastfeeding, before and after a frenulotomy, found abnormal tongue movements and ineffective removal of breast milk, followed by an improvement in milk transfer and tongue movement during breastfeeding after the tongue-clearing procedure.

We also observed a greater electric activity of the muscle in infants with a thin lingual fraenulum and with the frenulum attachment to the middle third of the tongue and visible from the sublingual caruncles, which had a sucking rhythm comprised of several sucks and short pauses, balanced coordination between food efficiency and sucking, swallowing and breathing functions, showing no signs of stress, and absence of bite of the nipple and tongue snapping, with a statistically significant difference when compared to the muscular electrical activity of infants with the fraenulum attached of the middle third of the tongue, between the middle third and the apex, and to the apex and visible from the lower alveolar ridge. A previous study [14] that analysed 109 full-term infants, and referred to a surgical procedure for tongue release (frenotomy), 14 of the infants who presented altered lingual fraenulum, showed a statistically significant relationship between anatomical characteristics of the lingual fraenulum and nutritive sucking, with improvement of the sucking pattern in breastfeeding after the frenotomy. Another previous study [41] identified that infants with an altered fraenulum were more likely to have difficulty sucking. Furthermore, a previous longitudinal study [15], which analysed the anatomical characteristics of the lingual fraenulum throughout the first year of the infant, observed that there is no change in its thickness, attachment of the tongue, and the floor of the mouth. Thus, early identification of the alteration is fundamental to intervention of the frenulotomy.

Previous studies have shown that the tongue actively participates in sucking and is essential for proper removal of breast milk [4,6–9]. The nipple is compressed from the tip to the base when the tongue is high and, when it lowers, the nipple expands by approaching the hard palate with the soft palate and increases in diameter causing a vacuum and allowing the milk to flow into the intraoral space [6–9]. Alterations of the lingual fraenulum may limit its motility [6,10–15], making it difficult the inadequate catching and can lead to changes in the sucking function, especially in the dynamic of sucking/removal of milk. The main problems identified in cases of altered lingual fraenulum in relation to breastfeeding in the mother’s womb are difficulties in catching, nipple pain and cracking, prolonged feeding time, reduced milk intake by the infant, loss of weight [11,14,33–35], dehydration, and growth deficiency [33–35]. Such changes may hinder the continuity of breastfeeding [10,11,14,33–35] with consequences for the infant’s health and, later on, the development of chewing and speech [10,13–15].

No studies associating electrical activity of the suprahyoid muscles and the lingual fraenulum of infants during feeding were found in literature. A previous study [5,20,22,24] that analysed the suprahyoid muscles during feeding using surface electromyography differed from the present study as to the objectives, infants’ age, sample size, feeding types and, because they did not associate the anatomical characteristics of the lingual fraenulum, made it impossible to compare the findings.

Surface electromyography has been shown to be effective in the early diagnosis of the limitations of tongue movements caused by the lingual fraenulum, whose attachment location, both on the tongue and the floor of the mouth, may raise doubts about the reduction of tongue mobility. This technique allows for the identification of possible oral motor dysfunctions, enables direct therapeutic interventions and early intervention, and consequently prevents feeding and communication alterations.

Among the main limitations of our study are the reduced schedules for the examination of the infants due to technical problems related to the regulation of the patients by the SUS, as well as the non-attendance of scheduled infants due to health problems and the public transportation strike.

Further studies investigating the the electrical activity of the suprahyoid muscles through surface electromyography after surgery to release the lingual fraenulum may contribute to a better understanding of the impact of ankyloglossia on tongue movements during breastfeeding.

## Conclusion

A greater electrical activity of the muscle was observed during sucking of the maternal breast in full-term infants when the lingual fraenulum was attached of the middle third of the tongue and visible from the sublingual caruncle. Lower electrical activity of the suprahyoid muscles during sucking of the maternal breast was observed in full-term infants with the lingual fraenulum attached at the apex of the tongue and visible from the lower alveolar ridge.

We also observed a greater electric activity of the muscle in infants with a thin lingual fraenulum and a rhythm comprised of several sucks and short pauses, adequate and balanced coordination between the feeding efficiency and the sucking, swallowing and breathing functions, without signs of stress, and absence of “bite” of the nipple and tongue snapping.

## Supporting information

Fig 1 - Electromyographic evaluation. Electrodes attached to bone (forehead) and submandibular (supra-hyoid muscles) region during sucking on breastfeeding

## Authors’ contributions

ECLF: conceptualization, data curation, formal analysis, writing original draft, writing reviw and editing. LACV: formal analysis, writing reviw and editing. IMFG: formal analysis, writing reviw and editing. RLCM: formal analysis, writing reviw and editing. CBS: conceptualization, data curation, formal analysis, writing original draft, writing reviw and editing. MAB: conceptualization, data curation, formal analysis, writing original draft, writing reviw and editing. All authors contributed to the revision and agreed upon the final article.

## Acknowledgement

We thank the employees of the Reference Center for Hearing Health of the Pontifical Catholic University of Goiás, and the mothers and newborns who participated in the study.

This study was funded by the Foundation for Research Support of the State of Goiás (Fundação de Amparo â Pesquisa do Estado de Goiás - FAPEG process: 2016/10267000710).

